# When will hybridization bring long-term benefits?

**DOI:** 10.1101/2025.01.16.633403

**Authors:** Hilde Schneemann, John J. Welch

## Abstract

Hybridization between distinct populations injects genetic variation, which can bring fitness benefits. However, these benefits often appear as F1 heterosis, and might not persist into later generations; especially since, as emphasized by classical theories, heterosis can be caused in several different ways. Here, we study the long-term outcomes of hybridization, using a model that allows us to tune several properties of the genetic variation, including the strength and architecture of heterosis, thereby unifying the classical theories. Results suggest that long-term outcomes depend mainly on the variance in epistasis, which determines the ruggedness of the fitness landscape, but without affecting the heterosis. Together, results suggest that the study of heterosis may tell us relatively little about the long-term outcomes of hybridization, and that hybridization might bring benefits more often than has been assumed.

## Introduction

When divergent populations meet and mate, the consequences for the hybridizing populations can vary. Sometimes the hybridization is beneficial, e.g. by establishing new, adaptive trait combinations, or increasing adaptive potential (Anderson, 1948; Chan et al., 2019; Kulmuni et al., 2023); but sometimes the hybridization is deleterious, e.g. by establishing maladaptive trait combinations, or swamping local adaptation (Rhymer and Simberloff, 1996). Predicting these diverse outcomes is an important goal, not only for understanding wild hybridizations (Arnold, 1992; Abbott et al., 2013; Peñalba et al., 2024), but also for plant and animal breeders (East, 1908; Shull, 1908; Gowen, 1952; Gerdes et al., 1999; Gur and Zamir, 2004; Mackay et al., 2020), and for conservationists considering translocations between isolated populations, or interventions against anthropogenic hybrids (Genovart, 2008; Aitken and Whitlock, 2013; Frankham, 2015; Chan et al., 2019).

To predict the outcome of hybridization, it is important to distinguish between its consequences in the short and long terms (East, 1936; Anderson, 1948; Gallais, 1988; Erickson and Fenster, 2006; Johnson et al., 2010; Schumer et al., 2016; Thompson and Schluter, 2022). First-generation F1 hybrids are often fitter than their parents – a phenomenon known as F1 heterosis (Darwin, 1876; Gowen, 1952; Lippman and Zamir, 2007; Mackay et al., 2020; Thompson and Schluter, 2022). But F1 are fully heterozygous and cannot breed true. So this early benefit of hybridization need not persist into later generations (Vetukhiv, 1954; Birchler, 2003; Koltunow and Tucker, 2003; Mackay et al., 2020).

Whether F1 heterosis does imply long-term potential benefit must depend on how it is caused (Crow, 1948; Koltunow and Tucker, 2003; Lippman and Zamir, 2007; De Sanctis et al., 2023). For example, under the classical “dominance theory”, heterosis is explained by parents having fixed recessive deleterious mutations, e.g. via inbreeding, population bottlenecks, or range expansion (Oakley et al., 2015, 2019; MacPherson et al., 2022). This explains heterosis via the masking of recessive effects in the F1 (Bruce, 1910; Keeble and Pellew, 1910; Crow, 1948); but it also implies that the deleterious mutations might be selected out in the longer term, leaving a high-fitness homozygous hybrid, containing the ancestral alleles.

By contrast, F1 heterosis would not imply long-term benefits if it were caused by overdominance – i.e. an intrinsic fitness benefit of heterozygosity (East, 1908; Shull, 1908; Crow, 1948), or if the parents contain coadapted gene complexes – i.e. sets of alleles that function well together, but poorly when only a subset of the alleles are present (Hill, 1982; Lynch, 1991; Simon et al., 2018). For this reason, several methods exist for investigating the causes of F1 heterosis, using e.g. comparisons of early generation hybrids (Cockerham, 1980; Hill, 1982; Lynch, 1991; Lynch and Walsh, 1998; Birchler, 2003; Lippman and Zamir, 2007; Wang et al., 2012; Jiang et al., 2017, e.g.), or introgressed genomic segments (see e.g., Wang et al., 2012; Torgeman and Zamir, 2023); however, it is not always clear what, exactly, we might infer from these data about the long-term potential of a hybridization (Lippman and Zamir, 2007).

Here, we study the process of hybridization as a whole, to explore the links between the initial process of divergence (Figure 1A), the strength and architecture of heterosis among early-generation hybrids (Figure 1B), and the long-term outcomes of hybridization (Figure 1C). Specifically, we perform individual-based simulations of secondary contact, but using a model where the genetic variation, which is introduced by the hybridization, can be tuned. This allows us to adjust the strength and architecture of heterosis, such as the relative contributions of epistasis or overdominance, and to investigate the influence of these factors – if any – on the long-term outcomes.

**Figure 1:**
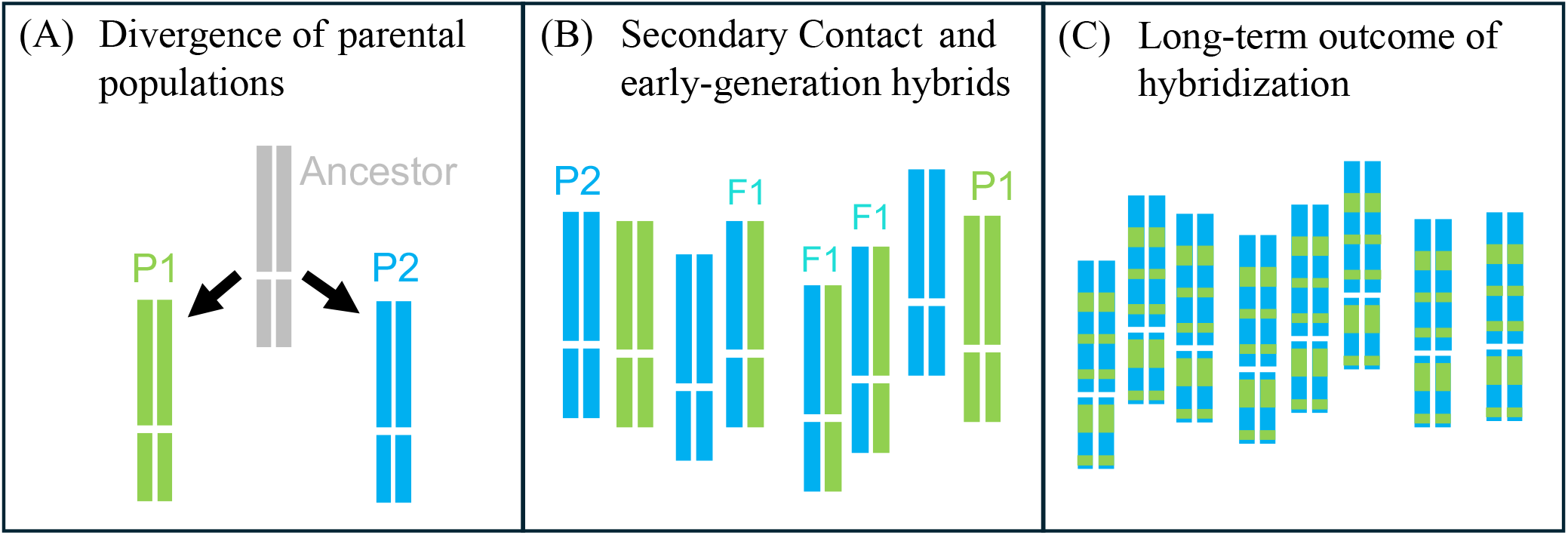
The scenario studied in this paper. **(A)** Two populations accrue genetic differences over a period of independent evolution. Divergence is via the fixation of random mutations, such as might occur under high inbreeding or low population size. **(B)** The diverged populations are then mixed in known proportions, injecting genetic variation. Initial F1 hybrids will tend to show heterosis, i.e. increased fitness. **(C)** The population continues to evolve as a hybrid swarm, with random mating, free recombination, selection and drift. Evolution ceases when the initial variation is exhausted, leaving a single genotype that is homozygous at all loci. This final genotype is the long-term outcome of hybridization, and its fitness can be compared to that of the ancestral genotype, the diverged parental lines, and the initial F1.

## Methods and Results

### Modelling parental divergence by random mutation accumulation

To model the divergence between diploid parental lines (Figure 1A), we followed the dominance theory (Crow, 1948), assuming that the parents accrue random mutations without effective selection. For reasons explained below, we implement this scenario using a simple phenotypic model, of optimizing selection on *n* quantitative traits (e.g. Lande, 1980; Turelli, 1985; Orr, 1998; Barton, 2001; De Sanctis et al., 2023). Under this model, which is illustrated in Figure 2A, the fitness of any genotype is determined from

**Figure 2:**
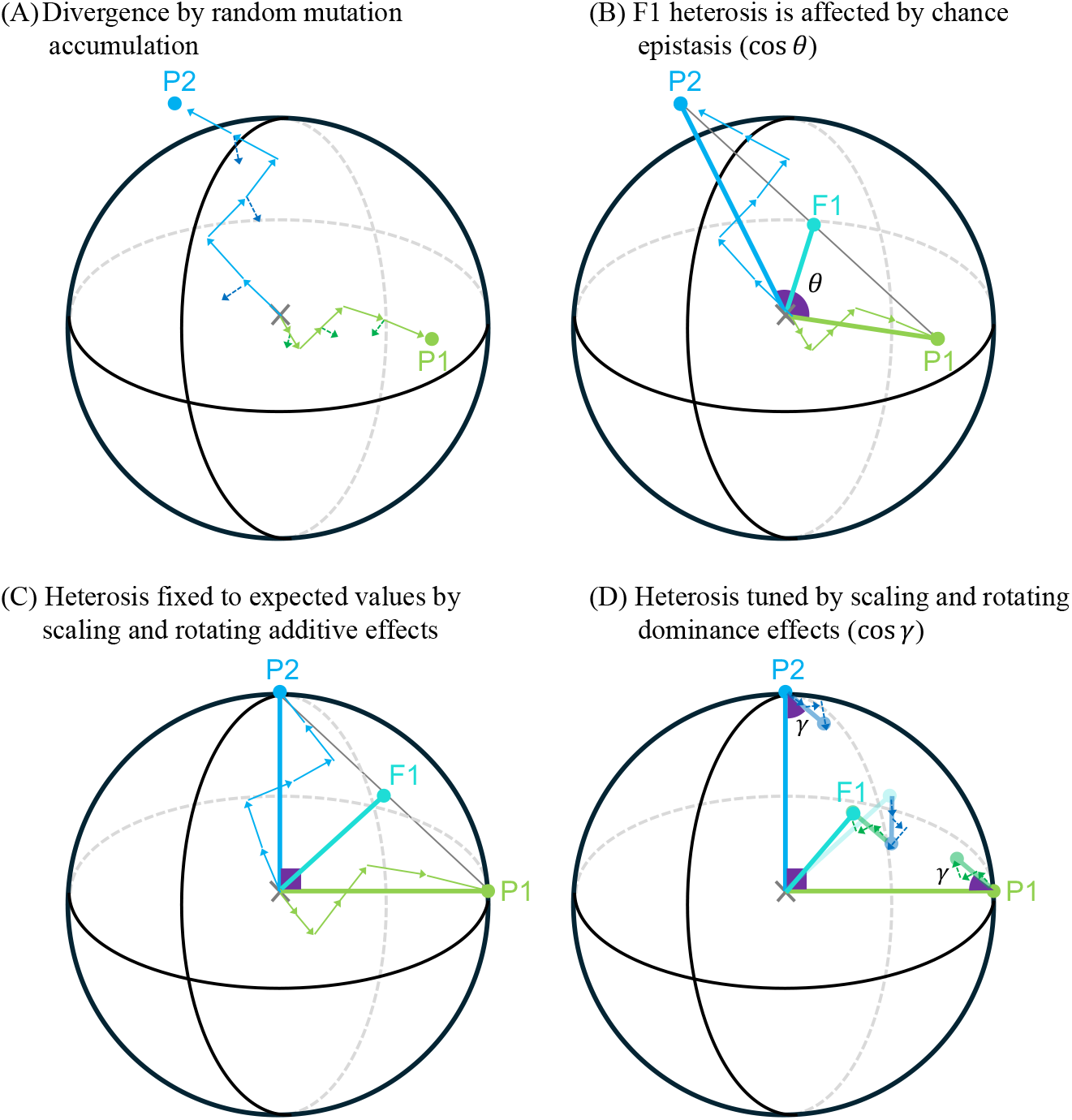
A model of random mutation accumulation, with tunable ruggedness in the fitness landscape, and tunable heterosis in the F1. Plots show a fitness landscape, which derives from a simple model of optimizing selection on *n* phenotypic traits (illustrated with *n* = 3). **(A)** Two parental lines, P1 and P2, begin at an optimal common ancestral state (*×*), and then accumulate *D* = 6 randomly-orientated mutations. Mutations, which are deleterious and recessive on average (eqs. 3-5), are characterized by an additive effect, **a**_*i*_ (lighter arrows), and a dominance effect, **d**_*i*_ (darker arrows) that contributes when the mutation is present in heterozygous state. The number of traits, *n*, affects the probability that random mutations act in similar phenotypic directions, and thereby the extent of fitness epistasis (eq. 7). **(B)** If we ignore the phenotypic dominance effects, the phenotype of the F1 hybrid matches the phenotypic midparent, and so its fitness depends on the fitnesses of the parents, and *θ*, the angle between them (eq. 10). *θ* also determines the contribution of epistasis to F1 heterosis (eq. 11). **(C)** By scaling and rotating the vectors of mutational effects, we can tune the strength and architecture of F1 heterosis – shown here, set at their expected values (eqs. 9 and 12). **(D)** We can also scale and rotate the phenotypic dominance effects, which affect F1 fitness by moving its phenotype away from the midparent. The important parameters here are *β* (the relative magnitude of the dominance effects) and *γ* (the relative orientations of the dominance and additive effects in the same parental line). When 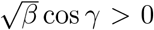, the dominance effects tend to compensate for the deleterious homozygous effects, which can increase F1 fitness (eq. 13).

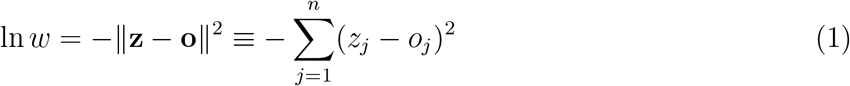

where **o** *≡ {o*_1_, *o*_2_, …, *o*_*n*_*}* is the optimal phenotype, and **z** *≡ {z*_1_, *z*_2_, …, *z*_*n*_*}* is the individual’s phenotype. To determine this phenotype, we associate a given mutation, *i*, with an additive effect **a**_*i*_ *≡ {a*_*i*1_, *a*_*i*2_, …, *a*_*in*_*}* and a dominance effect **d**_*i*_ *≡ {d*_*i*1_, *d*_*i*2_, …, *d*_*in*_*}*. The phenotype is the sum of these effects, distinguishing between mutations carried as homozygotes (Hom) or heterozygotes (Het):

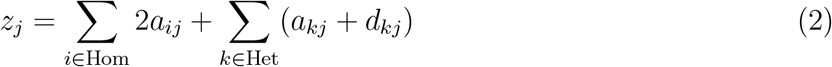

Given these assumptions, the fitness effects of new mutations depend on the sizes and orientations of their phenotypic effects. For example, in an optimally fit background, the selection coefficient of mutation *i*, is *s*_*i*_ ≈ *−*4∥**a**_*i*_∥^2^ for the homozygote, and *hs*_*i*_ ≈ *−*∥**a**_*i*_ + **d**_*i*_∥^2^ for the heterozygote (see Appendix S1.1 for full details and derivations). In the simulations reported below, additive effects were randomly orientated in phenotypic space, with a distribution of magnitudes that allowed us to control the strength of selection via a parameter 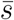.

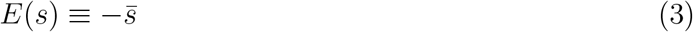

(see Appendix S1.1 for full details). Phenotypic dominance effects were generated in the same way, but scaled by a further parameter *β*, to control the relative sizes of the additive and dominance effects. It follows that *β* can affect the dominance coefficients of new mutations, depending on how this is estimated:

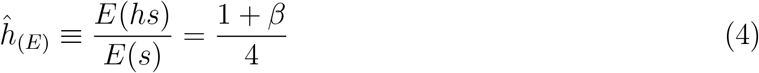

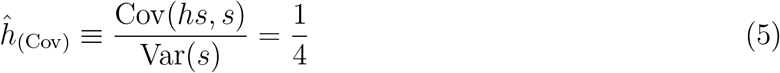

(see Manna et al., 2011; Schneemann et al., 2022 and Appendix S1.1). So with the first estimator (eq. 4), mutations are typically recessive if *β <* 1. With the second estimator (eq. 5), mutations are typically recessive, regardless of *β*. The assumption of recessivity is consistent with data (Manna et al., 2011; Billiard et al., 2021), and with the classical dominance theory of heterosis (Crow, 1948).

### Tuning the level of fitness interactions

In the phenotypic model described above, all fitness effects are context dependent. This allows us to extend classical single-locus theory (Crow, 1948), by including fitness interactions between loci (Martin et al., 2007; Fraïsse and Welch, 2019). The parameter *n* – the number of traits in the phenotypic model – is used to tune the extent of these fitness interactions. To see this, let us denote as *ε*_*ij*_ the pairwise epistatic effect between mutations *i* and *j*, defined as the deviation of their combined from the sum of their individual homozygous fitness effects: *ε*_*ij*_ = *s*_*ij*_ *− s*_*i*_ *− s*_*j*_ (i.e. the deviation of the double mutant from selective additivity). Summed across all pairs of divergent loci, this is equivalent to the additive-by-additive composite effect (Hill, 1982; Schneemann et al., 2024). This deviation can also be defined in terms of phenotypic effects as *ε*_*ij*_ *≈ −*8(**a**_*i*_ *·* **a**_*j*_) (Martin et al., 2007). It follows that mutations acting in random phenotypic directions will have zero epistasis on average, but will sometimes show negative or positive epistasis, by chance. Moreover, the variance in epistasis depends on *n*, as follows:

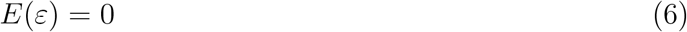

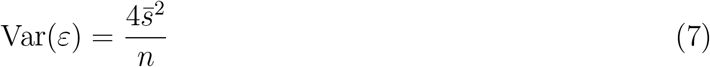

(see Welch and Waxman, 2003; Martin et al., 2007; Fraïsse and Welch, 2019, and Appendix S1.1 for details). So by varying *n*, we can alter the variance in epistatic effects between random mutations, while leaving other parameters unchanged. Varying *n* has two other effects, both related to the variance in epistasis, and both important to the results below.

First, let us note that any set of *D* mutations defines a fitness landscape comprising 2^*D*^ distinct haplotypes. Hwang et al. (2017) have shown that, when mutations are randomly orientated, the ratio *D/n* determines the ruggedness of this fitness landscape. So when *D/n* is small, it is very likely that the landscape will have a single fitness peak, meaning that only one of the possible genotypes has higher fitness than all of its single-mutant neighbors. By contrast, when *D/n* is large, there will be many such fitness peaks, and many combinations of alleles conferring relatively high fitness (Hwang et al., 2017).

Second, *n* affects the probability that a new randomly-directed mutation will point in the “right” direction, towards the optimum (Fisher, 1930). This, in turn, affects the ability of a population to maintain high fitness. For example, as shown by Lande (1980) and Barton (2016), under mutation-selection-drift balance at a stable optimum, the expected mean fitness decreases with the ratio *n/N*, where *N* is the population size:

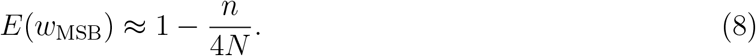

### Tuning the architecture of F1 heterosis

The phenotypic model also allows us to tune the strength and architecture of F1 heterosis, while leaving other quantities unchanged. To see this, let us consider the illustration of random mutation accumulation in Figure 2A. If the parents fixed a total of *D* mutations, with mean effect 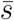, then the expected fitness of each line is simply

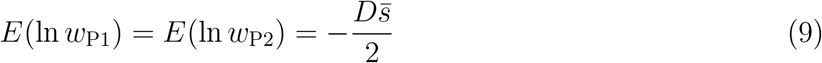

(e.g., Fraïsse and Welch, 2019). If we begin by ignoring the phenotypic dominance effects (assuming that *β* = 0) then as shown in Figure 2B, the F1 hybrid between these parents will match the midparental phenotype, and its log fitness follows from the cosine rule:

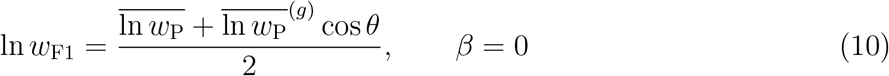

(Schneemann et al., 2022). Here, 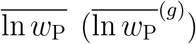 denotes the arithmetic (geometric) mean log fitness of the parents, and the second term depends on the angle between their phenotypes (Figure 2B). This second term also describes the contribution of epistasis to F1 heterosis (Hill, 1982), and is, in fact, proportional to the sum of the epistatic effects between the *D* mutations:

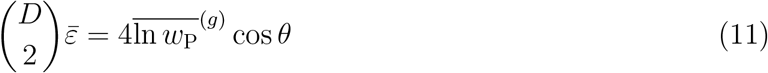

(see Chevin et al., 2014; Schneemann et al., 2020; De Sanctis et al., 2023 and Appendix S1.1 for details). In Figure 2A-B, positive epistasis arose from the (chance) tendency for the mutations in each parent to point in different phenotypic directions, thus compensating for the deleterious effects fixed in the other line. This led to cos *θ <* 0, and so to 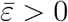.

If follows that we can tune the strength of heterosis, and the contribution of epistasis, by scaling and rotating the vectors of phenotypic change. For example, in Figure 2C we have manipulated a set of random mutations, so that the parents have exactly their expected fitness (eq. 9), and so does their F1:

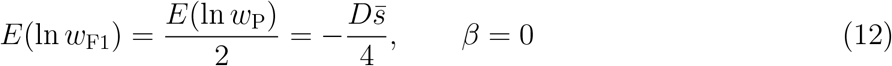

(Figure 2C). If we now consider the phenotypic dominance effects (Figure 2A), it is clear that their contribution to F1 heterosis can also be tuned by similar scaling and rotation. As illustrated in Figure 2D, phenotypic dominance effects move the F1 away from the midparent, and this may either decrease or increase F1 fitness (Schneemann et al., 2022). As such, an important quantity is the angle *γ*, which describes the tendency for dominance effects to compensate for additive effects, by pointing back toward the optimum. In the example shown in Figure 2D, F1 fitness is given by:

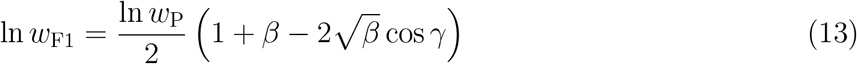

(see Appendix S1.1). So phenotypic dominance reduces F1 fitness if the effects are randomly orientated (i.e. if cos *γ* = 0; Schneemann et al., 2022), but can increase F1 fitness with compensation (i.e. if 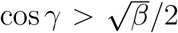). Note, too, that the model does not assume that compensation occurs at a single locus (so **d**_1_ is no more likely to compensate for **a**_1_ than for **a**_2_). This means that we can change *γ* without greatly altering the dominance coefficients of new mutations, such that eqs. 4-5 remain approximately unchanged (see Appendix S1.1). Biologically, however, our assumptions imply that compensation could only arise by chance, or if partially effective selection acts on the heterozygous effects of new mutations arising in already laden backgrounds (De Sanctis et al., 2023).

In summary, by simulating random mutation accumulation, with a given number of mutations (D) and strength of selection 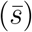, and then manipulating other parameters (*θ, β* and *γ*), we can control both the strength of F1 heterosis, and the relative contributions of dominance and epistasis to this heterosis (Hill, 1982). By changing *n*, we can independently control the ruggedness of the fitness landscape, via the variability of epistatic effects (Hwang et al., 2017).

### The long-term outcomes of hybridization

After modelling parental divergence, as described above, we simulated secondary contact and ongoing hybridization between the diverged parental populations (Figure 1B-C). The simulations assumed a simple Wright-Fisher population of a constant total size *N*. The initial *N* individuals contained a fixed proportion of P1 and P2 individuals, and since each line had accumulated *D/*2 random mutations, the hybrid population varied at *D* loci, which were assumed to be unlinked (see Discussion). Simulations included only selection and drift, and so there was no input from new mutations or new migrants. Simulations ended when all *D* loci had reached fixation for either the P1 or the P2 allele (Figure 1C; see also Appendix S1.3 for full details).

#### Long-term outcomes vary in single- versus multi-peaked fitness landscapes

Figure 3A shows the fitness of the final hybrid after replicated secondary contacts (Figure 1C), keeping several important parameters constant. In particular, we fixed the fitness of the hybridizing parental lines (ln *w*_P_), and of their initial F1 (ln *w*_F1_) – both set to their expected values under random mutation accumulation with additive phenotypes (see eq. 12 and Figure 2C). We also fixed the fitness of the ancestral genotype (ln *w*_Anc_ = ln *w*_Opt_ = 0), and the ratio *n/N* (thereby fixing the expected fitness at mutation selection balance: ln *w*_MSB_; eq. 8).

**Figure 3:**
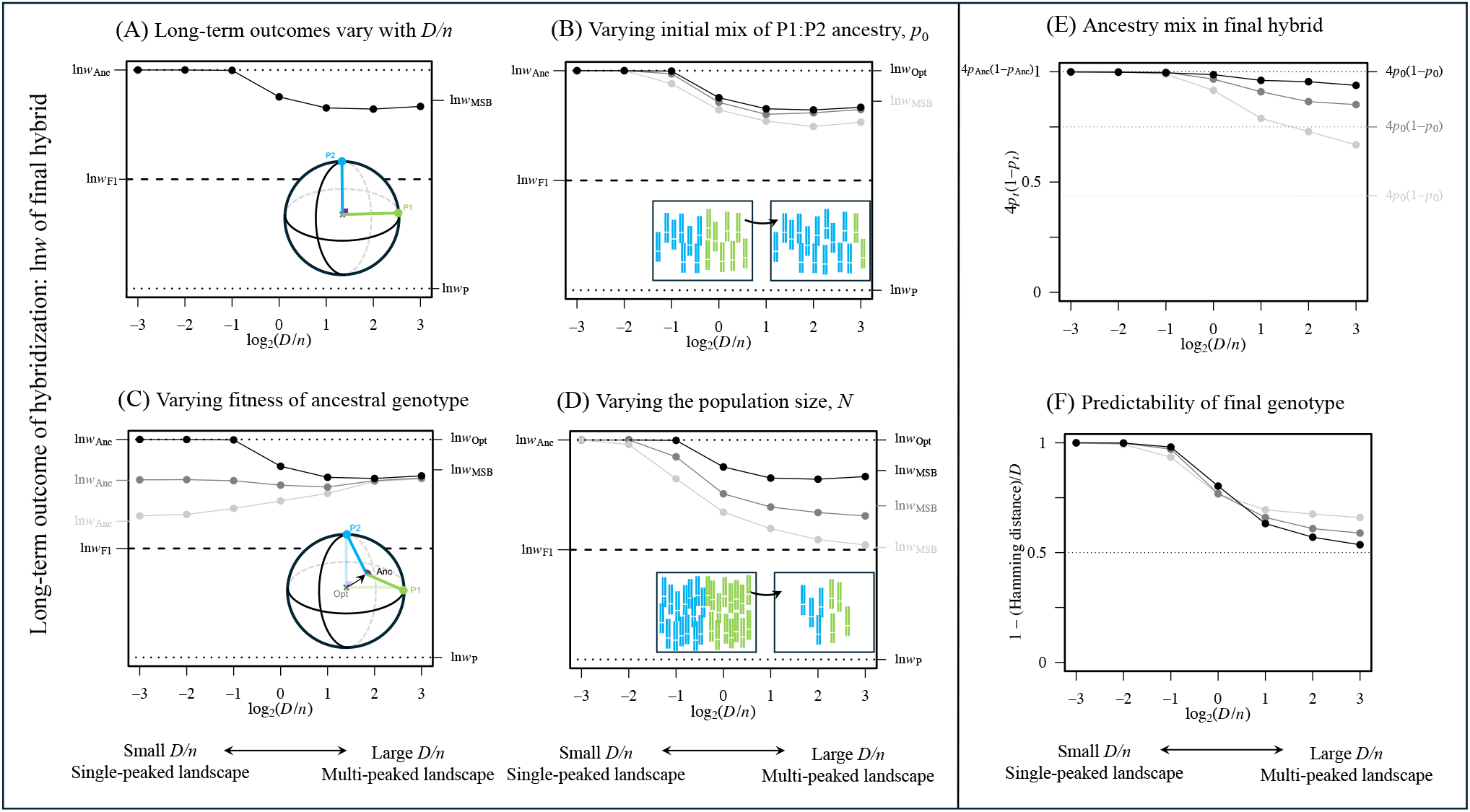
The ruggedness of the fitness landscape influences the long-term outcomes of hybridization. **(A)** Points show the log fitness of the final hybrid genotype, after simulated secondary contact (Figure 1C). Results are compared to the log fitnesses of the hybridizing parents (ln 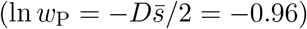), their initial F1 hybrid (ln 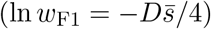), their common ancestor (ln *w*_Anc_), the optimal fitness (ln *w*_Opt_ = 0), and the expected fitness under mutation-selection balance: ln *w*_MSB_ = ln(1*−n/*4*N*) = ln(7*/*8). Outcomes depend on the ratio *D/n*, which determines whether the fitness landscape has one or many fitness peaks. **(B)** outcomes are relatively robust to varying the mix of ancestry present at secondary contact. (dark-to-light: proportion of P2 ancestry is *p*_0_ = 1*/*2, *p*_0_ = 1*/*4 and *p*_0_ = 1*/*8). **(C)** The fitness of the common ancestral genotype alters results when *D/n* is small (dark-to-light: ln *w*_Anc_ = 0, *−*0.18 and *−*0.36). **(D)** The population size alters results when *D/n* is large (dark-to-light: *n/*4*N* = 1*/*8, 2*/*8 and 3*/*8). The predictability of outcomes can also be measured by **(E)**: the proportion of P2 ancestry in the final hybrid (*p*_*t*_); or **(F)**: the mean number of sites that carry identical alleles after replicated secondary contact between the same parental lines. When *D/n* is small, hybridization predictably recovers the ancestral genotype (which had 50:50 ancestry); when *D/n* is large, the final genotype better reflects the mix of ancestry at secondary contact, but varies unpredictably at each locus. Each point is the median ((A)-(D)) or mean ((E)-(F)), over three sets of replicates with *D* = 24, 48 and 96; results were indistinguishable with the different *D* values as long as the key ratios were held constant.

Figure 3A does vary *D/n* – the parameter which determines the ruggedness of the fitness landscape (Hwang et al., 2017) – and results show a transition between two clear regimes. First, when *D/n* is small (left-hand-side of the plots), the single fitness peak corresponds to the ancestral genotype, before any random mutations had fixed. This means that, as long as selection against individual mutations is effective 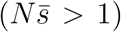, the long-term outcome of hybridization is always to recover this ancestral state. This is confirmed by the optimal fitness of the final hybrids (Figure 3B), and their 50:50 ancestry (Figure 3E), but also by the recovery of identical genotypes for replicated secondary contacts (Figure 3F). The importance of the ancestral state is confirmed in Figure 3C. Here, we moved the optimum, so that the ancestral state had suboptimal fitness, while leaving the other key quantities unchanged. In this case, when *D/n* was small, hybridization still led predictably to the ancestral genotype.

Now let us consider results when *D/n* is large (right-hand-side of plots). In this case, the fitness of the final hybrid matches expectations under mutation-selection-drift balance, and so is unaffected by the ancestral fitness (Figure 3C), but strongly affected by the population size (Figure 3D; eq. 8). In this regime, the mix of ancestry in the final hybrid better reflects the mix at secondary contact – i.e. if P2 individuals were initially rare, the final hybrid will contain less P2 ancestry, albeit with some increase of initially rare alleles (Figure 3E). Finally, the genotype of the final hybrid becomes unpredictable. Replicated secondary contacts can lead to similar fitnesses and similar mixtures of ancestry (Figure 3B and E), but to different alleles at any given locus (Figure 3F).

#### Long-term outcomes are little affected by the strength and architecture of heterosis

Figure 3 implies that long-term outcomes of hybridization could vary independently of the strength and architecture of F1 heterosis. This is confirmed in Figure 4. Figure 4A varies the strength of heterosis in the simplest way, by keeping the divergence model of Figures 2C and 3, but varying the fitness of the parents. As shown, as long as parents are sufficiently unfit (i.e. no fitter than ln *w*_Anc_ or ln *w*_MSB_), then the long-term benefits of hybridization are little affected. Figure 4B introduces phenotypic dominance effects, so that F1 fitness can be varied independently of the parental fitnesses (Figure 2D; eq. 13). Figure 4B contains results for the two extreme cases of no heterosis (*β* = 1, cos *γ* = 0, such that ln *w*_F1_ = ln *w*_P_), and maximal heterosis (*β* = 1, cos *γ* = 1, such that ln *w*_F1_ = ln *w*_Opt_). Nevertheless, the long-term outcomes of hybridization were little changed in either case (Figure 4B).

**Figure 4:**
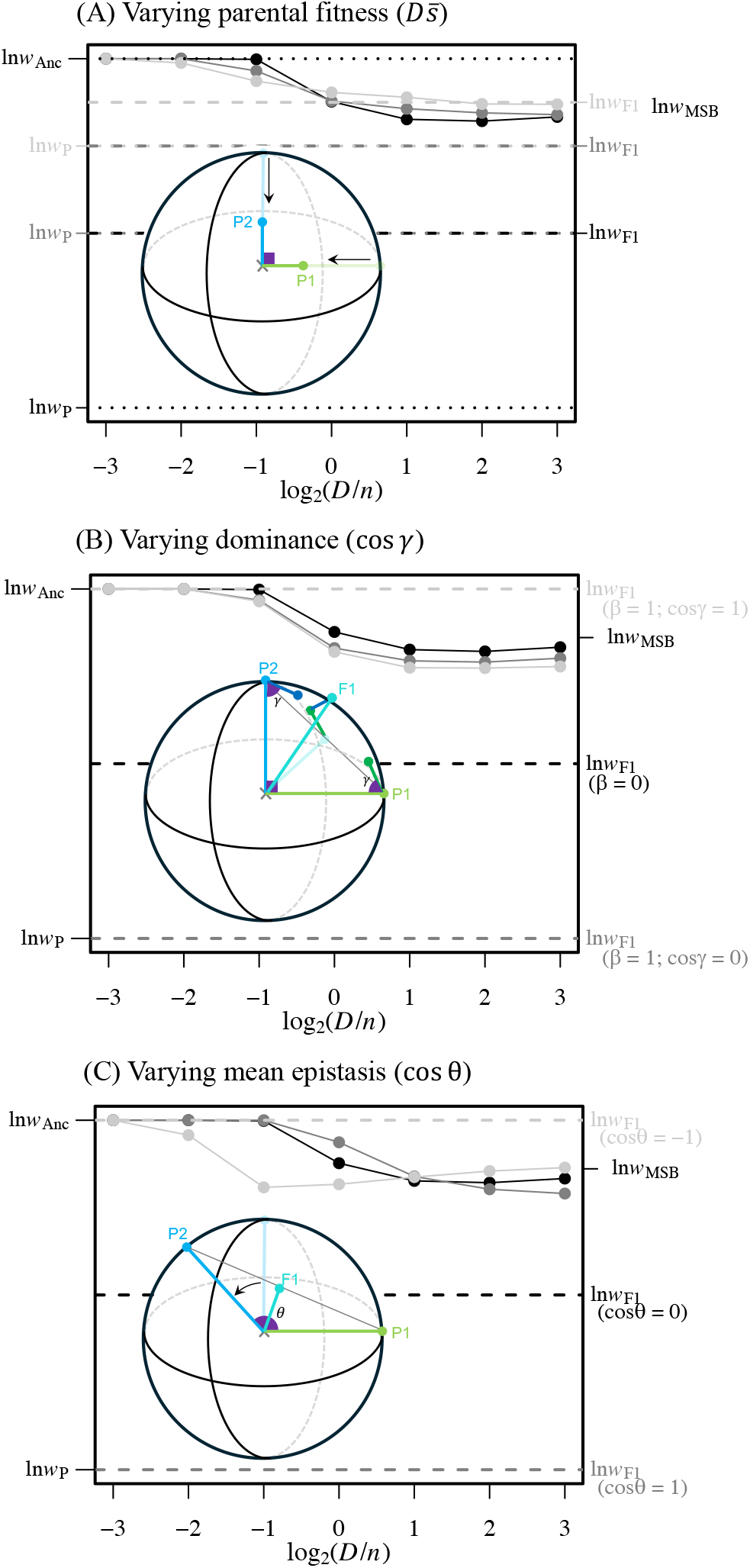
The strength and architecture of heterosis have little effect on long-term outcomes. All plots show the long-term outcomes of hybridization and match Figure 3A, except that F1 fitness is tuned by varying **(A)** parental fitness, **(B)** phenotypic dominance effects, or **(C)** mean levels of fitness epistasis. **(A)** Dark-to-light: 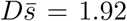, 0.96, and 0.48. **(B)**: Black: no phenotypic dominance (*β* = 0); Darker gray: randomly-orientated phenotypic dominance such that the F1 has no fitness advantage (*β* = 1, cos *γ* = 0); Lighter gray: dominance effects which tend to compensate for the additive effects, such that the F1 is optimally fit (*β* = 1, cos *γ* = 1). **(C)**: Black: no epistasis 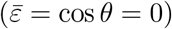; Darker gray: negative epistasis such that the F1 has no fitness advantage (*ε <* 0, cos *θ* = 1); Lighter gray: positive epistasis such that the F1 is optimally fit (*ε >* 0, cos *θ* = *−*1).

Finally, Figure 4C varies heterosis by introducing net levels of epistasis between the parental alleles. This was contrived by altering the angle *θ* between the parental phenotypes (Figure 2B; eqs. 10-11), and again, we considered the extreme cases of no heterosis (due to negative epistasis on average; cos *θ* = 1 such that 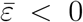) and maximial heterosis (due to positive epistasis on average; cos *θ* = *−*1, such that 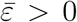). Results show that varying 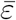 has qualitatively different effects in different parameter regimes. For example, when *D/n* is small, positive epistasis reduces the efficacy of selection against deleterious mutations (Kimura and Maruyama, 1966; Kondrashov, 1984), which can reduce the fitness of the final hybrid. By contrast, when *D/n* is large, positive epistasis increases the expected fitness of a random homozygous hybrid; this facilitates evolution in the multi-peaked landscape, increasing the fitness of the final hybrid. However, the latter effect is relatively small, while the former effect is strong only when *D/n ≈* 1.

### Random mutation accumulation can generate high levels of overdominance, but this has little effect on long-term outcomes

Although Figure 4 varied the architecture of heterosis, we did not explicitly consider overdominance (East, 1908; Shull, 1908; Crow, 1948; Lippman and Zamir, 2007). Overdominance can be difficult to measure, but one powerful approach uses introgression lines, where small regions of genome are introgressed from a donor parental line into a recipient parental line (Eshed and Zamir, 1994; Semel et al., 2006; Lippman and Zamir, 2007; Wang et al., 2012; Alseekh et al., 2013; Tian et al., 2019). As illustrated in Figure 5A-B, the effects of each introgression can be scored in both homozygous state (*s*_*I*_) and heterozygous state (*hs*_*I*_), and levels of overdominance assessed from the bivariate distribution of introgression effects. Figure 5B shows an empirical example, using published data from the genus *Solanum* (Semel et al., 2006; raw data kindly provided by Dr. Yaniv Semel; see Appendix S1.4 for full details). Introgressions in the blue areas of Figure 5B support predictions of the classical dominance theory (Crow, 1948), since they represent recessive deleterious introgressions (as expected if a recessive deleterious mutation had fixed in the donor line), and dominant beneficial introgressions (as expected if a recessive deleterious mutation had fixed in the recipient line, so that the introgression restored the ancestral allele). By contrast, introgressions falling in the red area of Figure 5B support the overdominance theory (Crow, 1948), since the heterozygous introgression outperforms both homozygotes.

**Figure 5:**
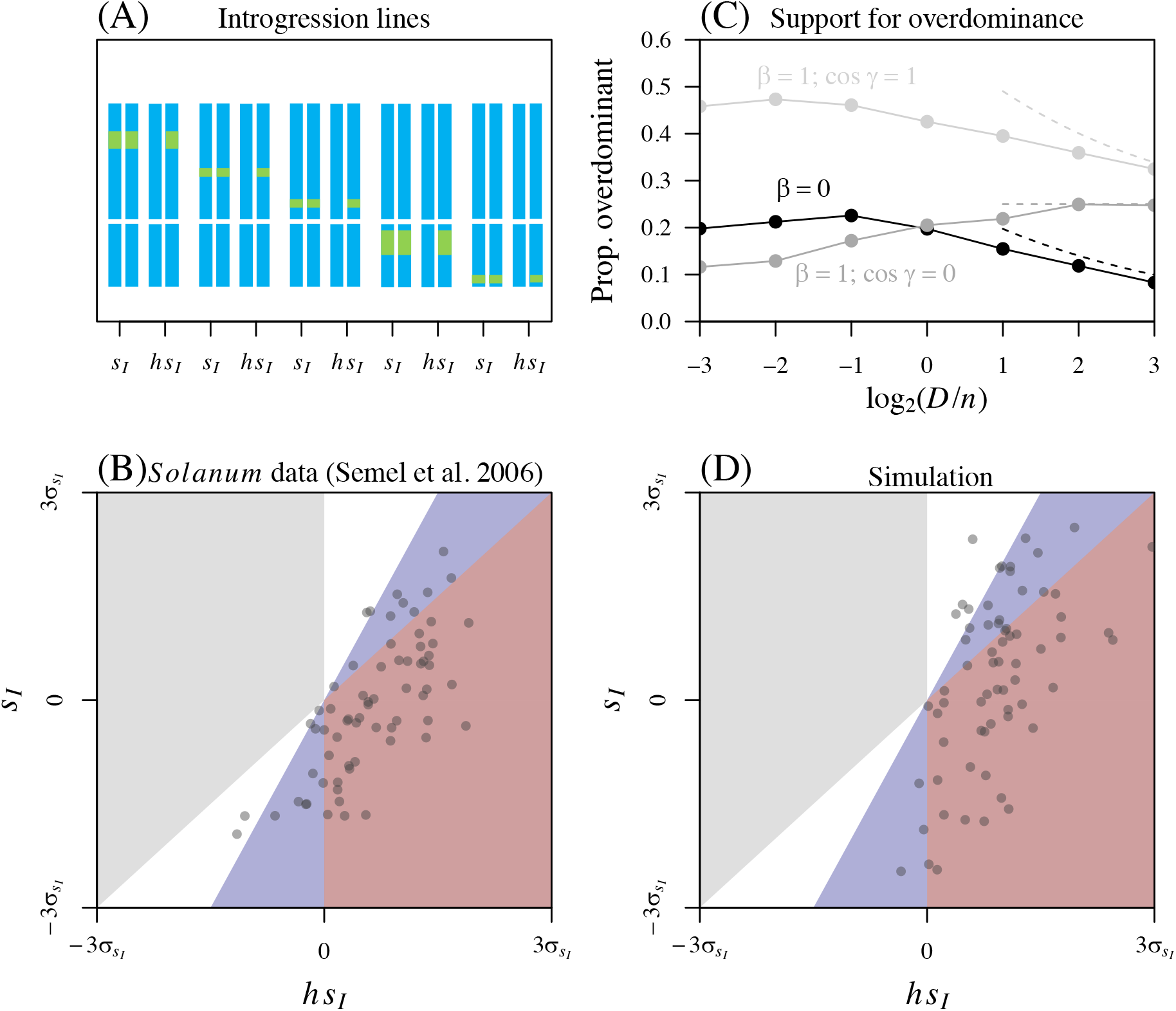
Distinguishing between dominance and overdominance using introgression lines. **(A)**: Introgression lines contain small regions of genome from a donor parental line introgressed into a recipient parental line (e.g. by repeated backcrossing). The effects of each introgression are measured in both homozygous and heterozygous state. **(B)**: Published data from 67 introgression lines between two species showing F1 heterosis, namely the wild tomato (*Solanum pennellii*; donor) and the cultivated tomato (*Solanum lycopersicum*; recipient) (Semel et al., 2006; and see Appendix S1.4 for details). Introgressions falling in the blue areas are deleterious recessive or beneficial dominant, and so support the dominance theory of heterosis. Introgressions falling in the red area are overdominant. Results are plotted in terms of the observed standard deviation of the homozygous effects: *σ*_*sI*_ = 0.1389. **(C)**: The proportion of simulated introgressions that are overdominant (falling in the red area of (B)), for the simulated parental lines reported in Figure 4B. Dashed lines show a bivariate normal prediction, extending results in Table 1, which holds when *D/n ≫* Var(*s*)/*s̄*^2^ (see Appendix S1.2 for details). **(D)**: a single set of simulated introgression lines, contrived to match the levels of overdominance in the *Solanum* data. We simulated *D* = 67 mutations, of which 50 fixed in the recipient line (matching the number of *Solanum* introgression lines, and the probable inbred state of the domesticated species). We then set *β* = 0.84, *D/n* = 1 and cos *θ* = cos *γ* = 1 to contrive the very high levels of overdominance.

Table 1 contains predictions for the bivariate distribution of introgression effects under our model of random mutation accumulation (see Appendix for derivation). It is notable that the results depend on all of the model parameters, including those accounting for fitness interactions; this is because introgressed alleles may interact with alleles present in the recipient background. The upshot is that our model – though derived under the assumptions of the classical dominance theory – can generate substantial levels of overdominance in introgression line data (see also Manna et al., 2011; Sellis et al., 2011). To see this, Figure 5C summarizes data from simulated introgression lines, which were generated from the parental lines used in Figure 4B. Comparing the two plots, it is clear that parameters such as *β* and *γ* can substantially affect the level of overdominance (Figure 5C) and yet have little or no effect on the long-term outcome of hybridization (Figure 4B). The only substantial effect of overdominance was to generate cases of persistent polymorphism.

**Table 1:** Predictions for introgression lines under random mutation accumulation.

|  | Introgression of derived allele | Introgression of ancestral allele |
| --- | --- | --- |
| $E(s_I)$ | $-\bar{s}(1 + 2\cos\theta)$ | $\bar{s}$ |
| $E(hs_I)$ | $-\bar{s}\left(\frac{1+\beta}{4} + \cos\theta(1 - \sqrt{\beta}\cos\gamma)\right)$ | $\bar{s}\left(\frac{3-\beta}{4} + \sqrt{\beta}\cos\gamma\right)$ |
| $\text{Var}(s_I)$ | $\text{Var}(s) + \frac{2D}{n}\bar{s}^2$ | $\text{Var}(s) + \frac{2D}{n}\bar{s}^2$ |
| $\text{Var}(hs_I)$ | $\frac{1+\beta^2}{16}\text{Var}(s) + \frac{1+\beta}{2}\frac{D}{n}\bar{s}^2$ | $\frac{9+\beta^2}{16}\text{Var}(s) + \frac{1+\beta}{2}\frac{D}{n}\bar{s}^2$ |
| $\text{Cov}(s_I, hs_I)$ | $\frac{1}{4}\text{Var}(s) + \frac{D}{n}\bar{s}^2$ | $\frac{3}{4}\text{Var}(s) + \frac{D}{n}\bar{s}^2$ |

However, this had a substantial effect on fixation times in only a small proportion of simulation runs, and never at a large number of loci (see also Ding and Goudet, 2005).

Even for simulations with the very high levels of overdominance observed in the *Solanum* data, long-term outcomes are as would be predicted from earlier results. For example, Figure 5D shows results from a simulation with *D* = *n* = 67, and with other parameters contrived to yield 45*/*67 overdominant introgressions, as compared to 44*/*67 for the real data. Nevertheless, replicated secondary contacts yielded the results consistent with our choice of *D/n* = 1, namely, a final fitness that was generally intermediate between the ancestral state, and mutation-selection-balance predictions (across 25 replicates with *N* = 1000: mean ln *w* = *−*0.015, range [*−*0.019, *−*0.010]; ln *w*_Anc_ = 0, ln *w*_MSB_ = ln(1 *−* 67*/*4000) *≈ −*0.017).

## Discussion

### What determines the long-term outcomes of hybridization?

This paper has explored the long-term outcomes of hybridization between genetically divergent populations. While hybridization always injects novel genetic variation, this variation can vary in both quantity and quality. Moreover, the short-term effects of hybridization – such as beneficial F1 heterosis – need not persist into the longer term, once heterozygosity has been lost (Koltunow and Tucker, 2003; Birchler, 2003; Mackay et al., 2020).

We have found that a major determinant of long-term outcomes is the overall amount of ruggedness in the fitness landscape, i.e. whether there are one or many fitness peaks. This is, in turn, determined by the variability – as opposed to the average – of epistatic interactions between alleles, and is governed in our model by the parameter *D/n* (Hwang et al., 2017; see Figure 3). When *D/n* is low, there is a single fitness peak and little variability in epistatic effects. In this case, deleterious mutations can be purged and the ancestral genotype recovered. By contrast, when *D/n* is large, this implies a rugged multi-peaked fitness landscape where the effects of alleles vary greatly with their genomic background. In this case, the final fitness of the hybrid population depends on the population size, consistent with predictions under mutation-selection-balance (Barton, 2016). The final homozygous genotype is no longer predictable, but its composition reflects the initial contribution of the two parental populations (Figure 3E-F). Note that, we assumed free recombination, so that *D* is the number of divergent sites, but in the general case with linkage, the relevant *D* is the number of linkage blocks, with the total effect of each block given by the joint effect of the mutations it carries. This implies that, with linkage, the relevant *D* may increase over the course of the hybridization, as recombination breaks up the initial linkage blocks, and that the final outcome will depend on a *D* that is somewhere between its two extremes of the number of segregating sites, and the number of chromosomes.

By contrast, and surprisingly, we found that long-term outcomes of hybridization are little affected either by the strength of F1 heterosis, or by its genetic architecture (see Figure 4). Results were little changed if the heterosis was caused by positive average epistasis, or by dominance effects (Hill, 1982); and also little changed by the amount of overdominance, as detected from introgression line data (see Figure 5). Here, the observed overdominance was not associative- or pseudo-overdominance, which is due to the linkage of mutations with independent effects (Jones, 1917; Collins, 1921; Lippman and Zamir, 2007). In our model, by contrast, single-locus heterozygotes could be truly advantageous, but only in certain backgrounds. This means that the overdominance is transient, coming and going as genetic backgrounds change (Sellis et al., 2011; Manna et al., 2011).

This hypothesized transient overdominance relies on fitness interactions between alleles, which are widespread in our model if *D/n >* 1 or *β >* 0. This assumption is consistent with data, including from mutation accumulation experiments (Halligan and Keightley, 2009; Crombie et al., 2024), and direct studies of effects in hybrid backgrounds (Wang et al., 2012; Jiang et al., 2017; Torgeman and Zamir, 2023; Frayer and Payseur, 2024), including context-dependent overdominance in rice hybrids (*Oryza sativa japonica × Oryza sativa indica*; Xie et al., 2022). Indirect support comes from the fact that species pairs with strong evidence of multilocus overdominance (Semel et al., 2006; Kaeppler, 2012; Tian et al., 2019), can nevertheless generate high fitness recombinant lines (Kaeppler, 2012). This is true of the *Solanum* species shown in Figure 5B (Christakis and Fasoulas, 2001; Gur and Zamir, 2015; Avdikos et al., 2021a,b). It is also true of the cotton species *Gossypium barbadense* and *Gossypium hirsutum*, which show high levels of overdominance in introgression lines (Tian et al., 2019), but have also undergone many successful introgression events in nature (Wang et al., 2022).

### Is hybridization always beneficial?

Another notable feature of our results is that hybridization led to a long-term fitness advantage in all cases considered. Indeed, the benefit of hybridization extended beyond the cases considered in the main text. For example, when the ancestral fitness (*w*_Anc_) and/or the mutation-selection-balance equilibrium (*w*_MSB_) are less fit than the original parental lines, hybridizing populations still find a combination of the parental alleles that increases fitness (see supplementary Figure S1). This may help to explain the widespread genomic signatures of hybridization in many taxa, even when early-generation hybrids are rarely observed (Mayr, 1963; Goodman et al., 1999; Mallet et al., 2007; Ottenburghs et al., 2015; Justyn et al., 2020; Kozak et al., 2021).

Of course, these results do not imply that hybridization will provide a fitness advantage in all possible cases. An obvious exception is when populations are locally adapted to different habitats – in which case hybridization can lead to allele swamping and loss of local adaptation (Rhymer and Simberloff, 1996; Wolf et al., 2001; Todesco et al., 2016). A second exception is when negative epistasis becomes so strong that all hybrids (or all post-F1 recombinant hybrids) have very low fitness. Negative epistasis is expected to increase steadily with divergence under a range of divergence scenarios (Hartl and Taubes, 1996; Schiffman and Ralph, 2021; Schneemann et al., 2020; De Sanctis et al., 2023; Schneemann et al., 2024). For example, under stabilizing selection, the expected F2 fitness is

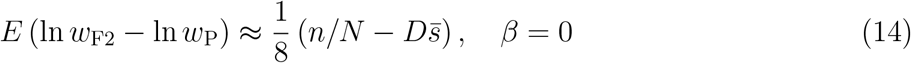

(see De Sanctis et al., 2023). So the segregation load grows with *D*, until reproductive isolation is essentially complete (Slatkin and Lande, 1994; Chevin et al., 2014; Simon et al., 2018).

Nevertheless, it is clear that both exceptions – local adaptation, and strong negative epistasis – are easily detectable over relatively short periods of time, e.g. by reciprocal transplant experiments, or from an F2 cross. And in the absence of these factors, the default expectation has to be that hybridization, with its injection of genetic variation, and potential increase in effective population size, will have positive fitness consequences in the long term (Schumer et al., 2018; Chan et al., 2019; Mackay et al., 2020; Kulmuni et al., 2023).

### A unifying theory of heterosis?

Even if heterosis does not help us to predict the long-term outcomes of hybridization, it remains a phenomenon of great interest (Gowen, 1952; Birchler, 2003; Kaeppler, 2012; Thompson and Schluter, 2022). Another notable feature of the present model, connected to its “tunability”, is that it generates heterosis in a wide variety of ways, despite our using a single, simple model of population divergence via random mutation accumulation. This mode of divergence is commonly associated with a particular theory of heterosis: the “dominance theory” (Crow, 1948). In line with this theory, heterosis arises primarily from the recessivity of deleterious mutations under our model (see eq. 4-5 and Manna et al., 2011), and we recover classical single-locus theory when fitness interactions are negligible (i.e., when *D/n ≪* 1 and *β* = 0). Nevertheless, we find that the strength of heterosis is also modulated by (chance) epistasis (eq. 32), and may be associated with background-dependent overdominance (see Table 1 and Figure 5), corresponding to other classical theories of heterosis (Crow, 1948; Hill, 1982). When these interactions are included (i.e., when *D/n >* 1 or *β >* 0), the same model generates data supporting several alternative theories (East, 1908; Shull, 1908; Keeble and Pellew, 1910; Bruce, 1910; Crow, 1948; Gowen, 1952; Lippman and Zamir, 2007).

The model is also highly flexible and has been used to study other scenarios, such as introgression between parents of very different fitness (Simon et al., 2018), heterosis caused by single locus overdominance (Manna et al., 2011), and bounded hybrid advantage, which varies with environmental conditions (Schneemann et al., 2020, 2022). All of these have been observed in nature (Vetukhiv and Beardmore, 1959; Moore, 1977; Eshed and Zamir, 1996; Wang et al., 1997; Arnold and Martin, 2010; Krieger et al., 2010; Sang et al., 2022; Nouhaud et al., 2022). As such, the model represents a significant step towards a unifying theory of heterosis (Birchler, 2003; Kaeppler, 2012).

## Acknowledgements

JW is very grateful to Nicolas Bierne, Ali Duncan and Guillaume Martin for their help with the LabEx CeMEB (Centre Méditerranéen de l’Environnement et de la Biodiversité) exchange program, for which this work was developed. HS acknowledges support from the Wellcome Trust program in Mathematical Genomics and Medicine (RG92770), and from the European Union’s Horizon 2020 research and innovation programme under the Marie Sk lodowska-Curie Grant Agreement No.101034413. Both authors are also grateful to the authors of the *Solanum* study for making their data publicly available, and especially to Dr. Yaniv Semel for providing the raw data. The authors distribute this work under a CC BY 4.0 copyright license.

## Appendix

### S1.1 Detailed methods for simulating random mutation accumulation

#### S1.1.1 Homozygous mutation effects

Let us consider a single new mutation *j* in homozygous form, and ask how its selection coefficient, *s*_*i*_, relates to its additive effects on the *n* phenotypes, **a**_*i*_ (eq. 2), via the fitness function of eq. 1. For simplicity, we will assume that the selective optimum is at the origin (**o** = **0**), and consistently use the following approximation, which applies when selective effects are small:

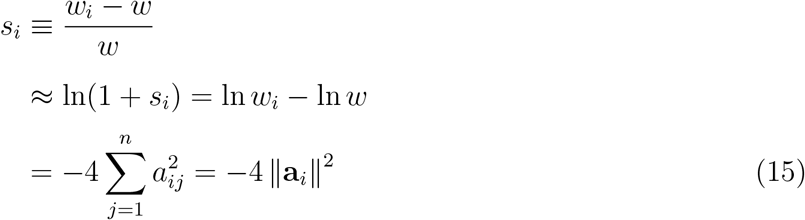

In our simulations, the mutation magnitudes, *∥***a**_*i*_*∥*, were drawn from a chi distribution with four degrees of freedom, and then multiplied by 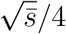.

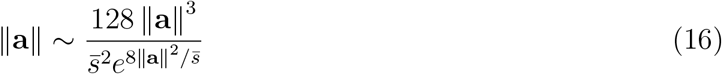

It follows from the moments of this distribution that the mean and variance in selection coefficients are:

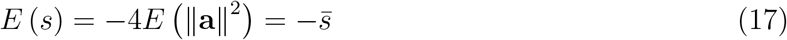

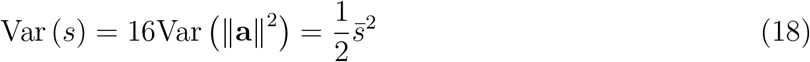

The fitness of each parental line depends on the sums of the additive effects of the *D/*2 mutations that they carry. If we order the mutations such that the first *D/*2 fixed in lineage P1, while the rest fixed in P2, then the fitnesses are given by

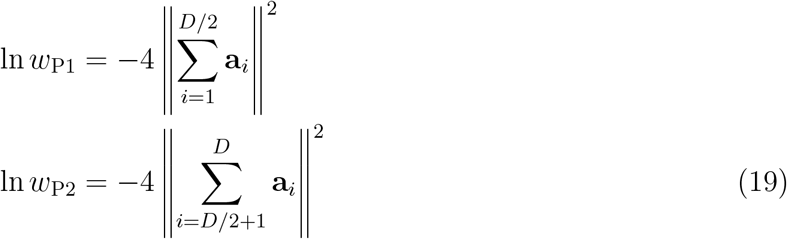

To control the parental fitness, we scaled the total length of the summed vectors such that it was equal to its expected value (eq. 9). In particular, we multiplied each magnitude by a factor such that

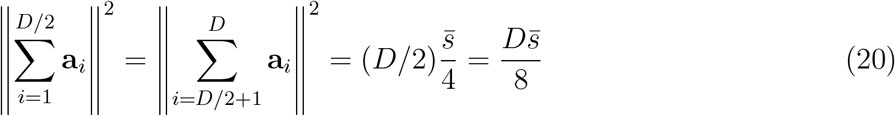

Note that this further scaling changes the variance in effect sizes (eq. 18), and also introduces non-independence between them. However, if *D* is sufficiently large, these effects will be small, and so are neglected in what follows.

##### S1.1.2 Epistatic interactions between homozygous mutations

The epistatic interaction between two homozygous mutations is defined via

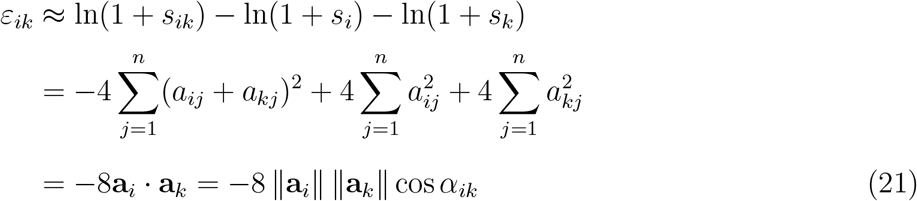

where *α*_*ik*_ is the angle in phenotypic space between the vectors of additive effects (Martin et al., 2007; Fraïsse and Welch, 2019). If the magnitudes are independent of each other, and of their orientations, then we have

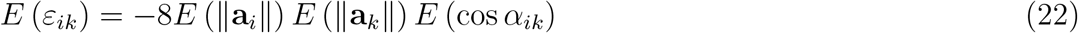

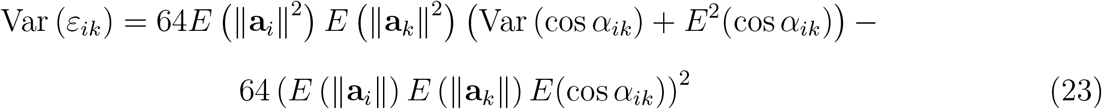

We assumed that mutations were randomly-orientated in phenotypic space (implemented by drawing their unscaled effects from a normal distrbution). In this case, the full distribution of cos *α* can be derived (see e.g., Welch and Waxman, 2003), but as shown by Fisher (1930), for sufficiently large *n*, it approaches a normal distribution with zero mean, and variance 1*/n*. Equations 6-7 then follow directly.

While we assumed no epistasis, on average, between the mutations fixed in each line, we did allow for new epistasis between the two sets of mutations. This was parameterized by the angle *θ* (Figure 2B), which was defined via the cosine similarity of the two sets of additive effects.

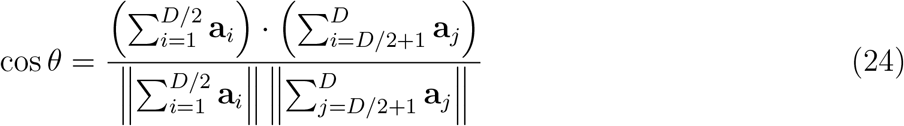

To see the relation to epistasis, consider the sum of epistatic effects between all *D* mutations that differentiate the parental lines.

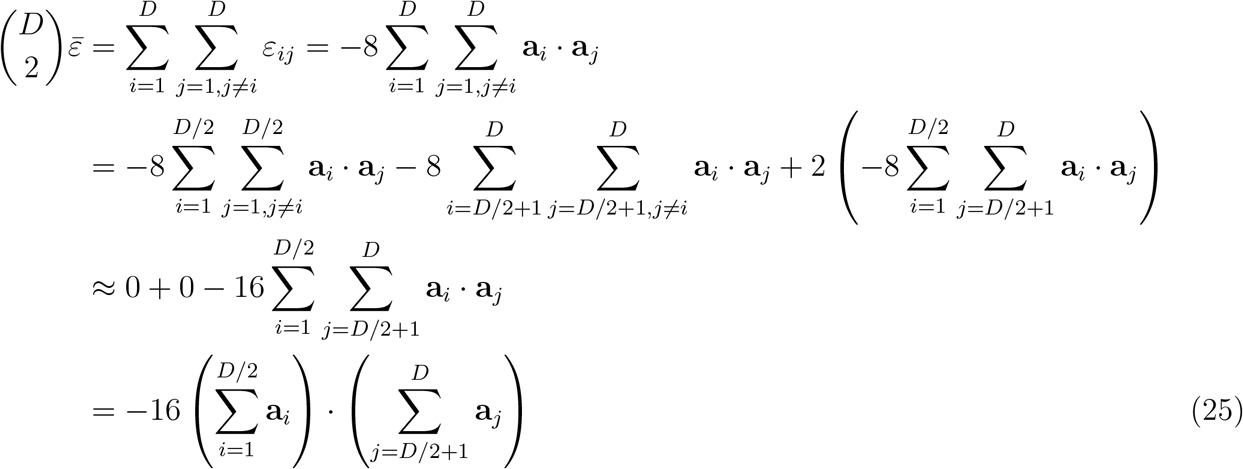

Combining this result with eqs. 19 and 24 yields eq. 11 of the main text. In the simulations, we fixed *θ* by rotating the vectors of summed additive effects in *n*-dimensional trait space, using an algorithm described by Zhelezov (2021).

##### S1.1.3 Heterozygous mutation effects

For the heterozygous effect of a mutation, we also need to consider its dominance effects on the *n* phenotypes, **d**_*i*_.

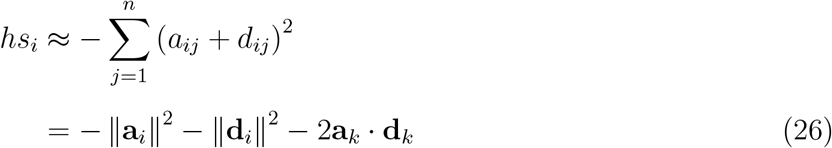

where 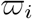 is the angle in *n*-dimensional phenotypic space between the additive and dominance effects of mutation *i*. In our simulations, the dominance effects were generated in the same way as the additive effects. In particular, they were randomly orientated, with magnitudes drawn from a scaled chi distribution, but in this case, also multiplied by a factor 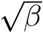.

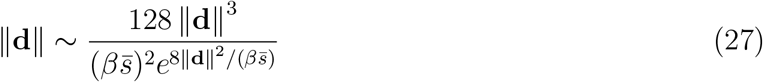

We also scaled the magnitude of the sum of dominance effects within each parental line, as described above, such that:

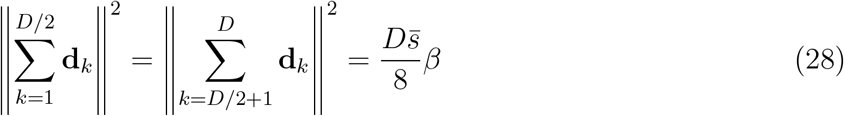

Finally, to tune the strength of heterosis using the dominance effects (eq. 13), we controlled the relative orientations of the summed additive and dominance effects. We did this via the angle *γ*, defined so that it took the same value for the sets of mutations fixed in P1 and P2:

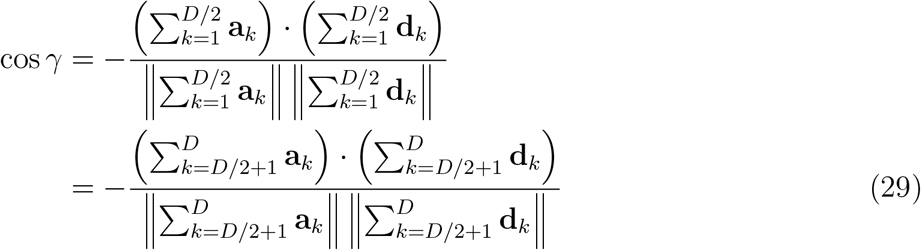

(see also Figure 2D). As described above, we fixed *γ* for each parental lineage, using vector rotation (Zhelezov, 2021).

The rotation of the dominance effects can also induce interactions between the dominance effects in one lineage, and the additive effects in the other. In particular, it follows from the procedures described above, that

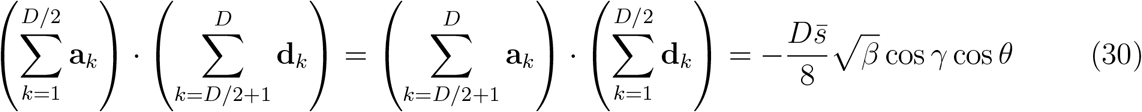

Rotation can also induce interactions between the two sets of dominance effects.

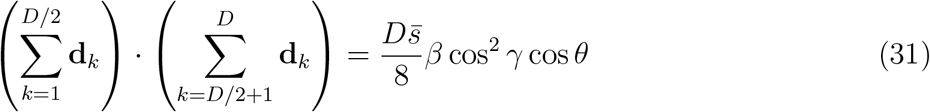

Putting all of these together, we can calculate the fitness of the F1 genotype.

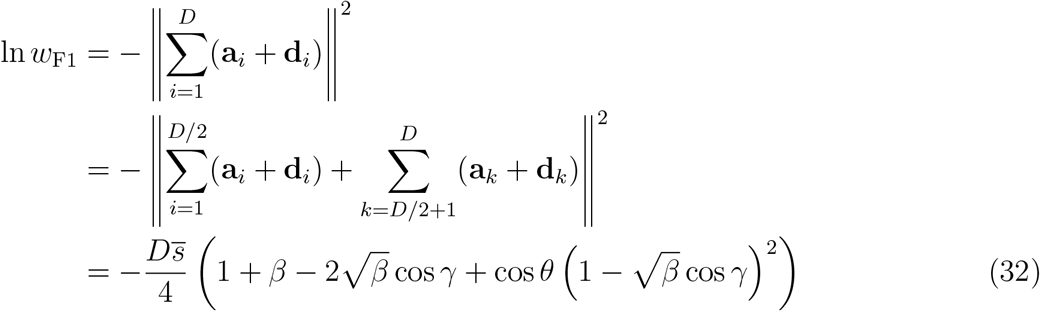

and equations 12 and 13 follow as special cases.

Let us now note that the rotation of the dominance effects, though designed to tune the overall strength of heterosis, also affects the properties of the individual mutations fixed in each line. However, because there was (by assumption) no special tendency for dominance effects at a given locus to compensate for the additive effects at the same locus, the effects on individual mutations were weak. To see this, note that using eqs. 20 and 28, we can write:

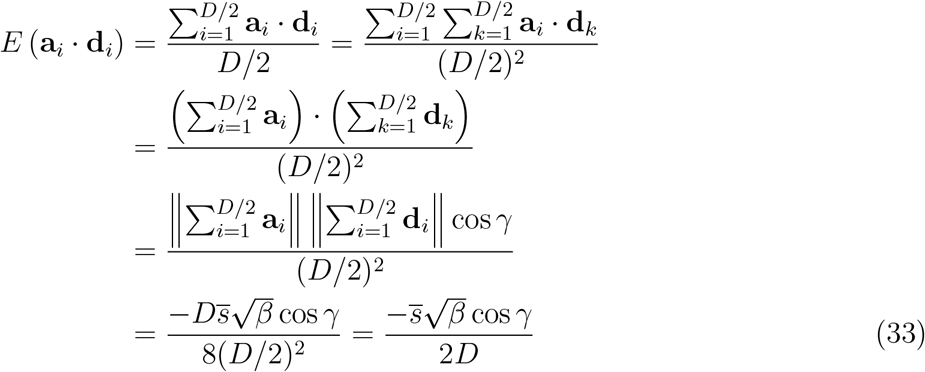

It follows that the mean heterozygous selection coefficient is

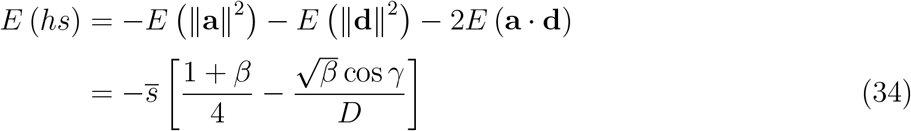

Main text equation 4 then follows, either as an approximation for large *D*, or exactly in the cases of *β* = 0 or cos *γ* = 0. The variance can be calculated in a similar way to eq. 23, i.e., by neglecting terms *𝒪* (*D*^*−*2^), such that *E*^2^(**a***·***d**) *≈* 0, and therefore Var(**a***·***d**) *≈ E* ∥**a**∥^2^ *E* ∥**d**∥^2^ */n*, as in eqs. 6-7. It follows that

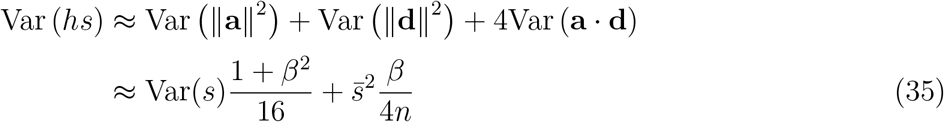

Similarly, for the covariance in heterozygous and homozygous effects, we have

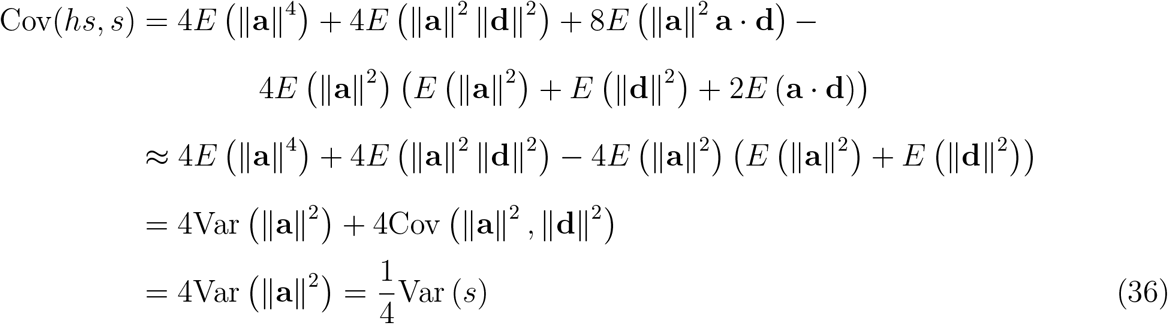

Equation 5 then follows directly.

#### S1.2 Predictions for Introgression lines

The results for introgression line data, shown in Table 1, follow the procedures outlined in section S1.1 above. If we treat P1 as the recipient line, then the effects of introgression *k* can be written

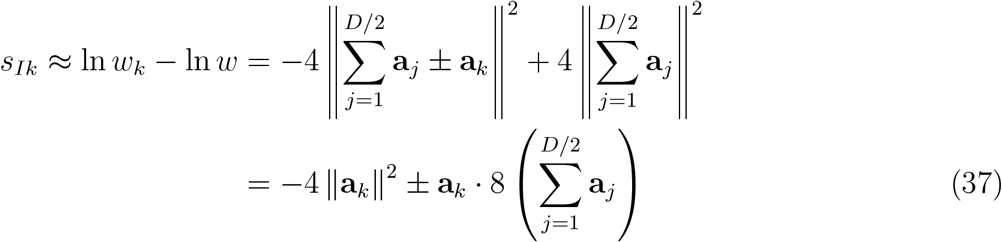

for the homozygote, and

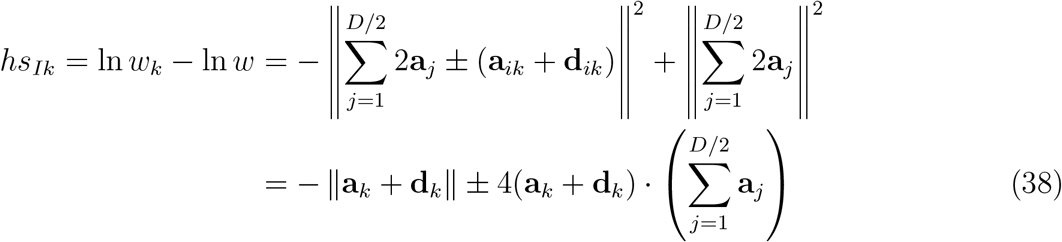

for the heterozygote. In both expressions, the plus-minus signs are positive for the introgression of a derived allele, which fixed in the donor P2 lineage such that *k > D/*2, and negative for the introgression of ancestral allele such that *k ≤ D/*2. It follows that, for the homozygous introgressions, we can write

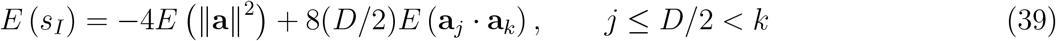

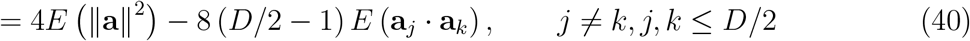

which apply to derived and ancestral introgressions respectively. Results in Table 1 then follow if we use the expectations for the intra- and inter-lineage interactions, derived in section S1.1 above (see, e.g., eqs. 6, 25, 29, 30, 31). The (co)variances, too, follow the procedures above, including the large-*D* approximations. The result is that all of the (co)variances in Table 1, involve two terms. The first term is proportional to the variance in single mutation effects, Var(*s*), and captures the direct effects of the introgressed allele; the second term is proportional to 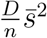, and captures the variance contributed by epistatic interactions (see eq. 6). If 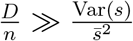 then we can neglect the first term, and the complete distribution of introgression effects (including both derived and ancestral introgressions), resembles a bivariate normal distribution, with covariance matrix 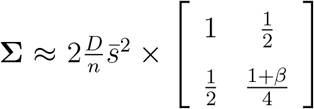. We can use this approximation to derive an expression for the proportion of introgressions that are overdominant when 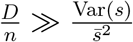.

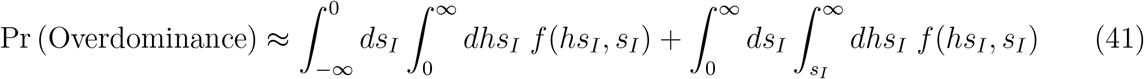

where *f* (*·, ·*) is a bivariate normal with the appropriate means and variances. Equation 41 can be solved numerically, or analytically for certain parameter combinations, including the cases shown by the dashed lines in Figure 5C.

#### S1.3 Simulations of secondary contact

To investigate secondary contact and ongoing hybridization, we used individual-based simulations of a Wright-Fisher population with selection. Simulations assumed a single hybrid swarm, with no spatial structure, and a constant-sized population of *N* individuals. Simulations were initiated with *Np*_0_ individuals of P2 ancestry and *N* (1 *− p*_0_) individuals of P1 ancestry. In each generation, 2*N* parents were chosen, with replacement, with probabilities proportional to their fitness, as determined from eqs. 1-2. Gametes were formed with free recombination at all *D* variable loci, and paired at random (i.e. with no assortative mating). For each combination of parameters, we simulated 5 divergence histories, each subject to 5 replicated secondary contacts (5*×* 5 replicates in all). This increased to 20 *×* 20 replicates for simulations with very small population size (*N <* 48). We ran simulations with three different values of *D* (*D* = 24, 48 and 96), but since results were indistinguishable for all three, plots show averages across the three sets of replicates. Other parameters were chosen to fix the key ratios, 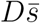, *D/n* and *n/N* as reported in the text and Figures. We chose these values to characterize the different regimes, but also so that *Np*_0_ was always an integer when *p*_0_ = 1*/*2. For this reason, some conditions were not run with smaller *D* when we set *p*_0_ = 1*/*4 and *p*_0_ = 1*/*8 (Figure 3B, E and F). While most simulation runs ended with fixation at all *D* loci, in a small number of cases, overdominance at a single locus led to persistent polymorphism (see main text). In these cases, simulations were terminated after 200*N* generations, and final fitnesses averaged across the remaining genotypes. We would expect such cases of balanced polymorphism to become more common in larger populations (with higher *N*), but they are not expected to affect many loci (Ding and Goudet, 2005).

#### S1.4 *Solanum* data curation

To make Figure 5B, we began with the raw data from Semel et al. (2006), including results from introgressions that did not meet the threshold to be counted as QTL. These were kindly provided by Dr. Yaniv Semel. We use seed weight per fruit as a proxy for fitness, since it was the trait with the most QTL, and because it correlates well with other reproductive traits in the data set (see Table 4 from Semel et al. 2006). We divided the trait values for seed weight per fruit by the maximum observed in this dataset. We then took the mean across individuals carrying a given introgression in homozygous or in heterozygous state, and subtracted the mean across 36 individuals of the recipient line M82. These are plotted as the values of *s*_*I*_ and *hs*_*I*_ in Figure 5B.

**Figure S1:**
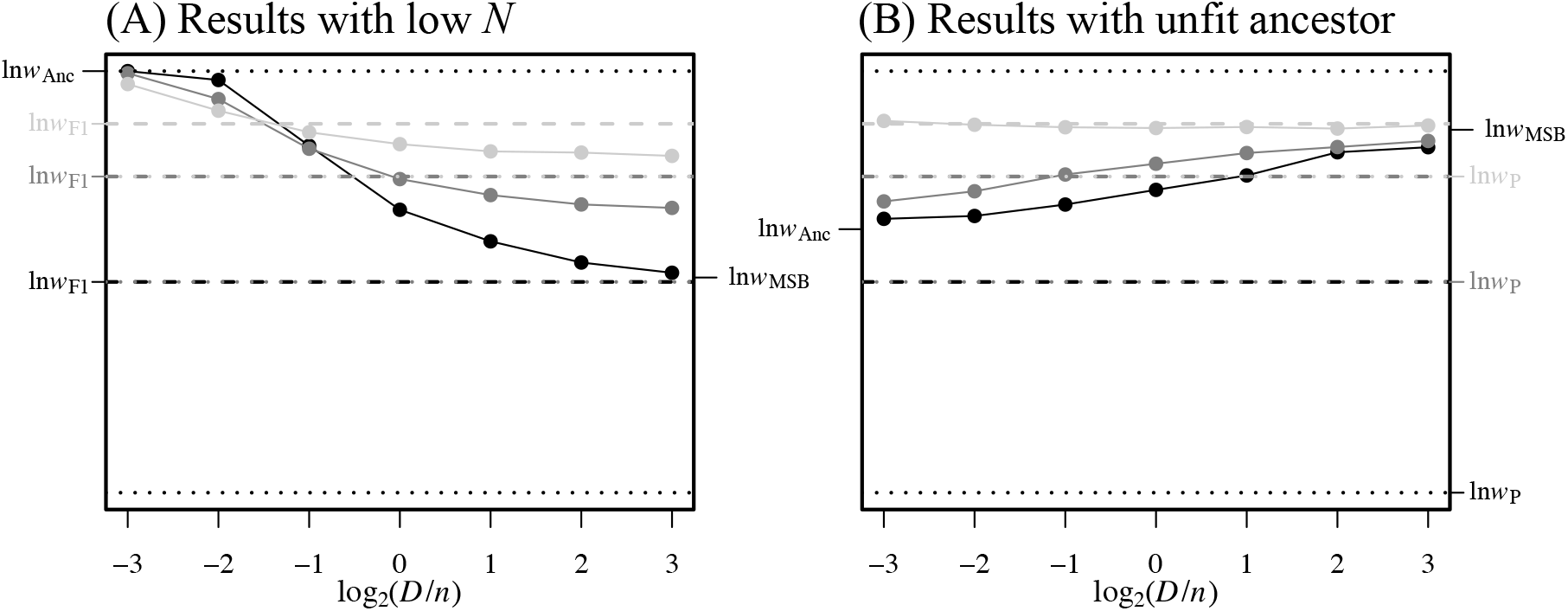
Hybridization can bring fitness advantages in additional cases. (**A**) If the parents are fitter than the the mutation-selection-balance equilibrium, then hybrids characterized by *D/n >* 1 may find a fitter combination of ancestral and derived alleles. (**B**) The same applies if *D/n <* 1, and the parents are already as fit or fitter than the ancestral state. All simulations and plotting details match Figure 3A.

